# Functional characterization of Rho GTPase activating proteins SYDE1 and SYDE2

**DOI:** 10.64898/2026.08.31.748432

**Authors:** Claire Song, Seong An, Hua Jane Lou, Benjamin E. Turk

## Abstract

The human genome encodes more than 60 proteins containing Rho GTPase activating protein (RhoGAP) domains, many of which remain understudied with respect to their target specificity and biological roles. SYDE1 and SYDE2 are two such orphan RhoGAPs, for which there are few studies characterizing their biochemical and cellular functions and conflicting reports identifying their cognate GTPases. We previously identified SYDE1 and SYDE2 in a screen for substrates of the c-Jun N-terminal kinases. Here, we show that SYDE1 and SYDE2 are preferentially phosphorylated by JNK1 relative to other mitogen-activated protein kinases (MAPKs) at sites proximal to a kinase docking region. Purified SYDE1 and SYDE2 are shown to have significant catalytic GAP activity toward RhoA, Rac1, and Cdc42. However, neither up-nor down-regulation of SYDE1/2 expression leads to detectable changes in bulk GTP loading of any of these GTPases. Nevertheless, we demonstrate that SYDE1 and SYDE2, in a partially GAP-dependent manner, increase cell spreading and number of focal adhesions, and promote more directionally persistent migration in HEK293 cells. Together, these findings establish SYDE1 and SYDE2 as robust JNK substrates with catalytic activity toward a set of Rho GTPases and reveal basic functions of SYDE1 and SYDE2 in regulating cell morphology, adhesion, and migration.

**One sentence summary:** A study of RhoGAPs SYDE1 and SYDE2 reveals their core biochemical and functional characteristics, and establishes their importance in cell morphology, adhesion, and migration.

## INTRODUCTION

The Rho family of small GTPases controls essential cellular processes related to cytoskeletal dynamics, morphology, adhesion, and migration ^1–3^. The family comprises 20 members, among which RhoA, Rac1, and Cdc42 are the best studied ^4–7^. Like all members of the Ras superfamily, Rho GTPases function as molecular switches that cycle between a GTP-bound “on” state, in which they bind to downstream effectors, and a GDP-bound “off” state. The nucleotide binding state is controlled by Rho guanine nucleotide exchange factors (GEFs) and Rho GTPase activating proteins (GAPs). GEFs catalyze exchange of GDP for GTP to promote the on state, while GAPs accelerate the intrinsic GTP-hydrolysis activity to promote the off state.

More than 60 human proteins containing RhoGAP domains have been described ^6,8^, far exceeding the number of Rho GTPases. Many RhoGAPs lack clearly defined Rho GTPase targets. Indeed, only a minority of RhoGAPs have been experimentally verified to have activity toward specific GTPases ^6,8^. Because many of these orphan GAPs are understudied in their catalytic and functional properties, their specific downstream signaling output and biological functions remain poorly understood. In-depth characterization of orphan GAPs at both the biochemical and cellular level is therefore needed.

The RhoGAP protein syd-1 was discovered through a genetic screen for regulators of synapse formation in *C. elegans* ^9^, and two mammalian homologs, SYDE1 and SYDE2, were subsequently identified. SYDE1 and SYDE2 contain a RhoGAP domain immediately downstream of a C2 domain, as well as long unstructured N- and C-terminal regions (**Fig. 1A**). The worm ortholog lacks a conserved Arg residue required by nearly all GAPs to accelerate GTP hydrolysis and appears inactive, although it can bind to and negatively regulate the Rho GTPase MIG-2 ^9,10^. The fly and mouse orthologs, by contrast, do harbor the catalytic Arg residue and reportedly act as GAPs toward Rac1/Cdc42 and RhoA, respectively ^11,12^.

**Fig. 1.**
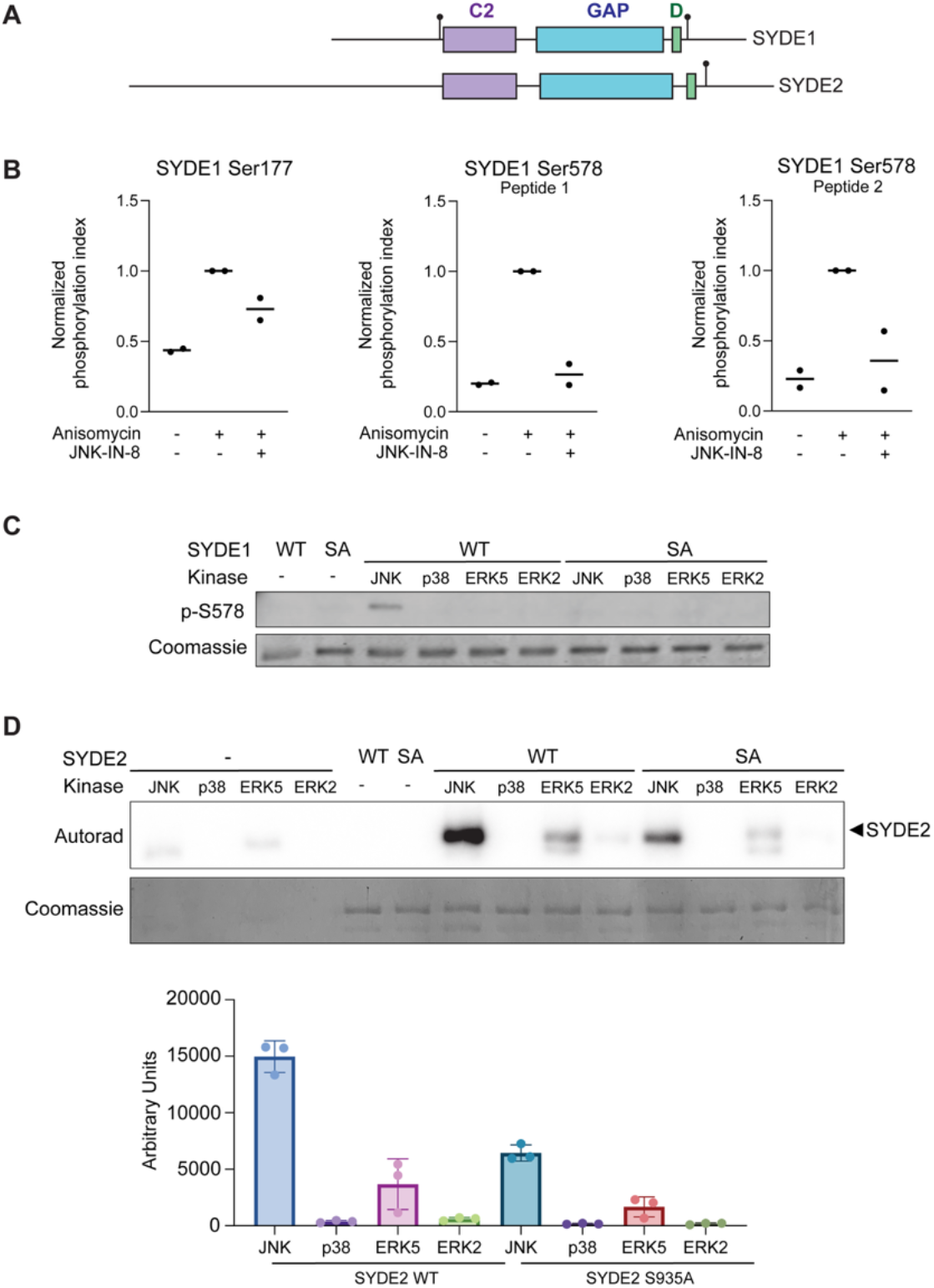
SYDE1 and SYDE2 are preferentially phosphorylated by JNK but are also phosphorylated by other MAPKs. (**A**) Domain schemes of SYDE1 and SYDE2 showing C2, GAP, and MAPK docking (D) regions. Black lollipops represent JNK-modulated Ser-Pro phosphorylation sites (SYDE1 Ser177, SYDE1 Ser578, and SYDE2 Ser935). (**B**) LC-MS/MS analysis of tryptic peptides from affinity purified SYDE1-3xFlag. SYDE1 was transiently expressed in HEK293T cells, which were treated as indicated. Graphs show ratios of the signal from the indicated phosphopeptide to that of the corresponding unphosphorylated peptide. Data are normalized to the anisomycin treated sample. Bars show mean of n = 2 biological replicates for Ser177, and n = 2 biological replicates for two distinct tryptic peptides phosphorylated at Ser578 (peptide 1, GGPE**<u>S</u>**PPSNR; peptide 2, GRGGPE**<u>S</u>**PPSNR). (**C**) In vitro kinase assay measuring MAPK-mediated phosphorylation of SYDE1 WT and S578A mutant (SA) with a phospho-specific SYDE1 Ser578 antibody (p-S578). A representative set of images from three separate experiments is shown. (**D**) In vitro radiolabel kinase assay measuring MAPK-mediated phosphorylation of SYDE2 WT and S935A mutant (SA). Representative images from three separate experiments is shown. Graph shows mean ± SD of quantified autoradiography signals from all three experiments.

Proteome-wide GAP surveys using direct and indirect activity readouts have yielded conflicting results with respect to the specific Rho GTPase targets of mammalian SYDE1 and SYDE2. A FRET-based biosensor screen for RhoA, Rac1, and Cdc42 activity suggested that SYDE1 targets RhoA and Rac1, while SYDE2 targets Rac1 alone ^13^. In contrast, a proximity labeling screen showed that SYDE1 can target Cdc42, RhoQ, and RhoJ ^14^. A separate siRNA screen in endothelial cells suggested that SYDE1 targets Rac1 and Cdc42 ^15^. This lack of consensus underscores the need for further investigation to determine relevant Rho GTPases for SYDE1 and SYDE2 in specific contexts.

The core biological functions of SYDE1 and SYDE2 in relevant cellular and organismal systems have also not been extensively characterized. Existing evidence suggests SYDE1 may play an important role in neuronal ^9,11,12,16,17^ and placental biology ^18,19^, although whether these roles require catalytic RhoGAP function is unclear. In the worm and fly, Syd-1 respectively regulates axon guidance and synaptic assembly ^9–11,16,17^, and mouse SYDE1 regulates synaptogenic signaling and vesicle docking ^12^. *Syde1* knockout in mice results in aberrant placental vascularization, indicating its potential importance outside of the nervous system ^18^. Fewer studies address SYDE1 and SYDE2 function in isolated cellular systems. In primary endothelial cells, SYDE1 reportedly decreases endothelial barrier integrity ^15^. *SYDE1* knockdown in human JAR trophoblastic cancer cells inhibits migration ^18^, and both SYDE1 and SYDE2 localize to focal adhesions when overexpressed in fibroblasts ^13^, suggesting a potential role in actin cytoskeleton-dependent phenotypes. Overall, a more complete functional characterization of SYDE1 and SYDE2 is needed to clarify their biological roles.

We previously identified SYDE1 and SYDE2 in a screen for c-Jun N-terminal kinase 1 (JNK1) interactors from a library of short peptide fragments of human proteins ^20^. Here, we further characterize the functions of SYDE1 and SYDE2, assessing their phosphorylation by multiple mitogen-activated protein kinases (MAPKs), their GAP activity against a panel of Rho GTPases in vitro and in cells, and their effects on cell morphology, focal adhesions, and migration. We found that both SYDE1 and SYDE2 are specifically phosphorylated by JNK1 over other MAPKs. We further show that both proteins exhibit significant GAP activity toward RhoA, Rac1, and Cdc42 in vitro. We reveal that SYDE1 and SYDE2 promote cell spreading in HEK293 cells in a manner partially dependent on catalytic GAP activity. The increase in cell spreading was associated with a higher number of focal adhesions and altered migration. Together, these results provide new biochemical and cellular insights into SYDE1 and SYDE2 function, further illuminating their biological roles.

## RESULTS

### SYDE1 and SYDE2 are specifically phosphorylated by JNK

We previously identified SYDE1 and SYDE2 in screening a library of short peptide fragments of human proteins for sequences binding to the JNK1 docking groove, a protein interaction site within the kinase domain separate from the catalytic cleft ^20^. The corresponding JNK docking sequences are located at an analogous position in SYDE1 and SYDE2 C-terminal to the RhoGAP domain (**Fig. 1A**). We further found that both SYDE1 and SYDE2 could serve as JNK1 substrates in vitro and in cells, and we mapped a JNK-dependent phosphorylation site on SYDE2 ^20^ (**Fig. 1A**). To identify potential JNK phosphorylation sites in SYDE1, we isolated transiently-expressed SYDE1 from HEK293T cells and conducted liquid chromatography-tandem mass spectrometry (LC-MS/MS) analysis. Cells expressing SYDE1 were treated with the protein synthesis inhibitor anisomycin to activate the JNK pathway in the presence or absence of a selective JNK inhibitor, JNK-IN-8 ^21^. We found that anisomycin induced phosphorylation at two residues, Ser177 and Ser578, which was blocked by JNK-IN-8 treatment (**Fig. 1A, B**).

Although MAP kinases are activated by distinct stimuli, all MAPKs phosphorylate substrates in the context of identical Ser/Thr-Pro motifs ^22^. Therefore, MAPKs other than JNK could theoretically phosphorylate SYDE1 and SYDE2 under non-stress conditions. To examine the capacity of different MAPKs to phosphorylate SYDE1 Ser578 and SYDE2 Ser935, we conducted in vitro kinase assays with representatives from the canonical MAPK subfamilies (JNK1, p38α, ERK5, and ERK2). We examined phosphorylation of SYDE1 Ser578 in particular because it also occupies a position analogous to SYDE2 Ser935, proximal to the JNK docking region (**Fig. 1A, B**). Because the specific activity of recombinant kinases can vary depending on the level of activating phosphorylation among other factors, we first assayed each kinase on a common MAPK peptide substrate, allowing us to include equivalent units of activity in SYDE kinase reactions. Using an antibody raised against SYDE1 phospho-Ser578, we found that this site was phosphorylated solely by JNK, with no detectable signal for the other MAPKs. A SYDE1 S578A mutant provided no signal for any of the kinase reactions, confirming phospho-specificity of the antibody (**Fig. 1C**). To assess SYDE2 phosphorylation, we conducted in vitro radiolabel kinase assays and found that it was also preferentially phosphorylated by JNK1, but it was additionally phosphorylated to a lesser extent by ERK5 and ERK2. SYDE2 S935A showed attenuated phosphorylation with all kinases compared to WT SYDE2 though did not completely eliminate phosphorylation by JNK1 and ERK5. These kinases therefore appear to phosphorylate other sites on SYDE2, though to a lesser extent than Ser935 (**Fig. 1D**). Taken together, these results confirm that SYDE1 Ser578 and SYDE2 Ser935 are specifically and robustly phosphorylated by JNK1 over other MAPKs.

### Human SYDE1 and mouse SYDE2 have GAP activity towards RhoA, Rac1, and Cdc42 in vitro

SYDE1 and SYDE2 harbor RhoGAP domains yet have not been assessed for catalytic GAP activity towards a panel of Rho-family GTPases. To determine whether SYDE1 or SYDE2 displayed GAP activity on RhoA, Rac1, and Cdc42, we conducted in vitro GTPase activity assays using purified human SYDE1 (residues 172-587) and mouse SYDE2 (residues 507-944) constructs harboring the C2 and GAP domains. We found that WT SYDE1 and SYDE2 had significant GAP activity towards RhoA, Rac1, and Cdc42, which was reduced by mutating the conserved catalytic Arg finger residue (SYDE1 R369A and SYDE2 R707A) (**Fig. 2A, B**). Phospho-regulation of GAP activity has been reported in vitro for other GAP proteins ^23–28^, and phosphorylation can directly alter catalytic GAP activity by affecting autoinhibition or homodimerization ^4,23,29,30^. We thus investigated whether JNK phosphorylation might impact the intrinsic activity of SYDE1 or SYDE2. Because we were unable to purify the phosphorylated form of SYDE1, we created a phosphomimetic substitution at Ser578 (S578E, SE). We found that the phosphomimetic version did not significantly change GAP activity towards Rac1 and Cdc42 compared to WT SYDE1, though it did appear to slightly increase GAP activity towards RhoA (**Fig. 2A**). To assess whether JNK phosphorylation of SYDE2 affects its catalytic GAP activity, we conducted GAP assays using phosphorylated and unphosphorylated full-length forms of SYDE2 WT and RA purified from HEK293T cells treated with anisomycin and JNK-IN-8 (**Fig. 2C**). We previously found that anisomycin treatment of ectopically expressed SYDE2 leads to near complete JNK-dependent phosphorylation at Ser935 ^20^. We confirmed the phosphorylation states of these proteins by electrophoretic mobility shift on Phos-tag SDS-polyacrylamide gel electrophoresis (PAGE). We found that phosphorylation of SYDE2 by JNK did not significantly affect catalytic GAP activity towards RhoA, Rac1, or Cdc42. Altogether, these results demonstrate that phosphorylation of SYDE1 or SYDE2 likely does not directly modulate their catalytic GAP activity in a significant manner, at least as measured in vitro. However, these experiments do not rule out a role for phosphorylation in modulating GAP activity indirectly by changing subcellular localization or intermolecular interactions.

**Fig. 2.**
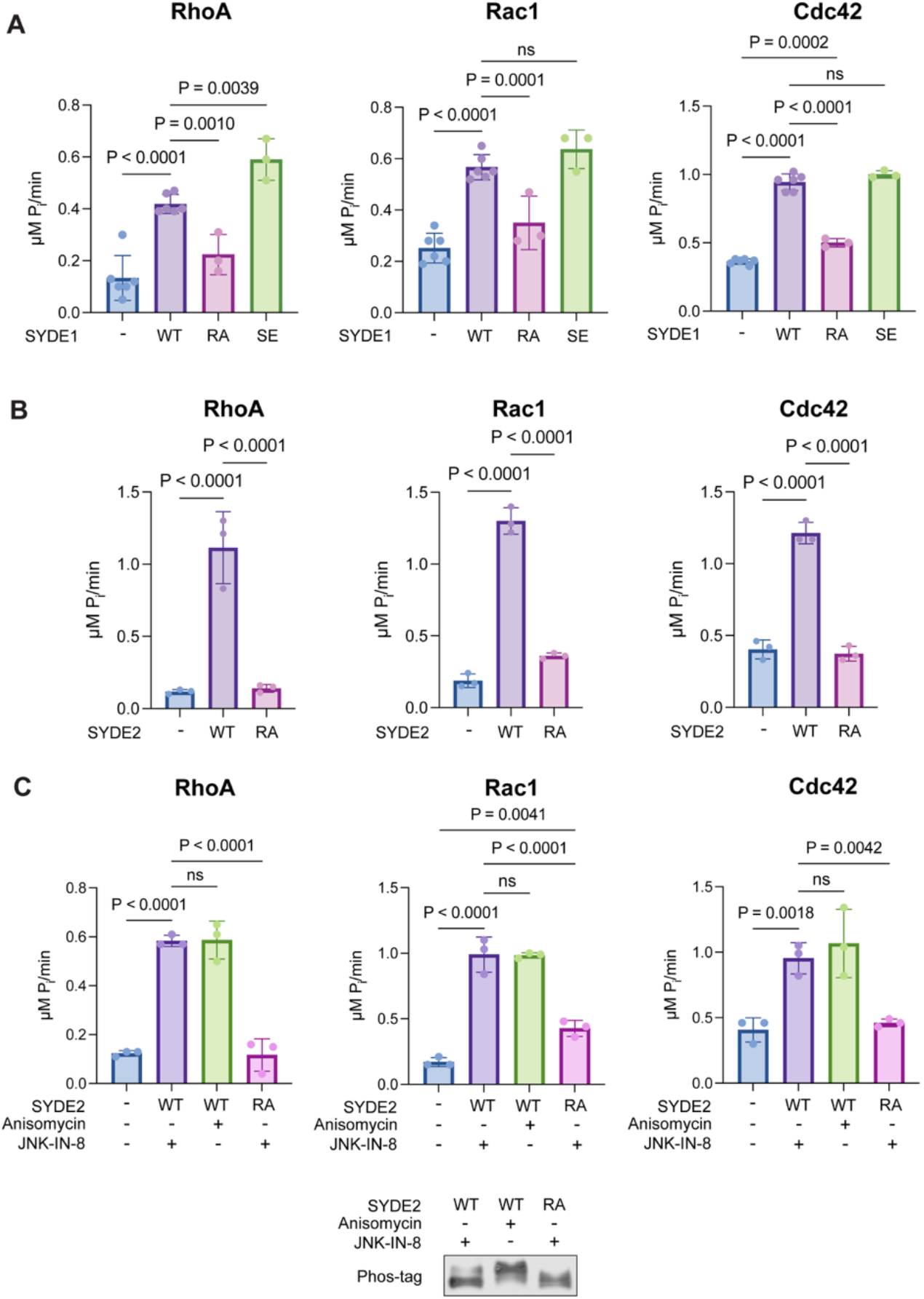
SYDE1 and SYDE2 have GAP activity towards RhoA, Rac1, and Cdc42 in vitro. (**A**) Multiple turnover GAP activity assays with 500 nM SYDE1 and 5 μM RhoA, Rac1, or Cdc42. WT, wildtype; RA, R369A GAP-inactive mutant; SE, S578E phosphomimetic mutant. Data shown as mean ± SD from independent replicates (n ≥ 3). P values were calculated using ordinary one-way ANOVA with Tukey’s multiple comparisons test. (**B**) Multiple turnover GAP activity assays with 25 nM SYDE2 and 5 μM of RhoA, Rac1, or Cdc42. WT, wildtype; RA, R707A GAP-inactive mutant. Data shown as mean ± SD from independent replicates (n = 3). P values were calculated as for panel (A). (**C**) Multiple turnover GAP activity assays with 25 nM unphosphorylated (isolated from cells treated with JNK-IN-8 alone) or phosphorylated (isolated from cell treated with anisomycin) SYDE2 and 5 μM of RhoA, Rac1, or Cdc42. SYDE2 variants and significance calculations are as in panel (B). Bottom panel shows Phos-tag SDS-PAGE analysis to confirm phosphorylation states of SYDE2 used in the assays.

### SYDE1 and SYDE2 do not significantly affect global GTP-loading of RhoA, Rac1, or Cdc42 in cells

Because SYDE1 and SYDE2 had significant catalytic GAP activity towards RhoA, Rac1, and Cdc42 in vitro, we investigated whether they reduced overall GTP loading of any of these GTPases in cells. As SYDE1 is thought to play an important role in cytoskeletal remodeling, cell migration, and differentiation in placental trophoblast cells ^18,19^, we initially conducted experiments in Swan 71 cells, a telomerase-immortalized first trimester trophoblast cell line that retains key characteristics of primary trophoblast cells^31^. We generated a clonal Swan 71 *SYDE1* knockout line by CRISPR/Cas9 gene editing and stably re-expressed WT or mutant forms (R369A, S578E) of SYDE1-GFP to close to endogenous levels (**Fig. S1**). Using this knockout/rescue system, we assessed overall levels of GTP-loaded RhoA, Rac1, and Cdc42 using G-LISA effector binding assays. We found no significant changes in GTP-loading of RhoA, Rac1, or Cdc42 upon re-expression of WT SYDE1 or the RA or SE mutants in SYDE1 KO Swan 71 cells (**Fig. 3A**). Because we did not see an effect at endogenous levels of SYDE1, we next examined whether ectopic overexpression of either SYDE1 or SYDE2 might impact GTP loading. We stably expressed SYDE1-GFP (WT and R369A, S578E, S578A mutants) or SYDE2-GFP (WT and R707A, S935D, S935A mutants) (**Fig. S1**) and similarly measured levels of GTP-loaded GTPase. We again found no significant changes to GTP-loading in HEK293 overexpressing any form of SYDE1 or SYDE2 (**Fig. 3B, C**). Although SYDE1 and SYDE2 have catalytic GAP activity in vitro towards RhoA, Rac1, and Cdc42, it is possible that in a cellular context low-level or spatially-restricted activation of the GTPases may contribute to undetectable changes in bulk GTP-loading by the G-LISA readout. Alternatively, as GAPs can be promiscuous in vitro, SYDE1 and SYDE2 may target other GTPases in a cellular environment.

**Fig. 3.**
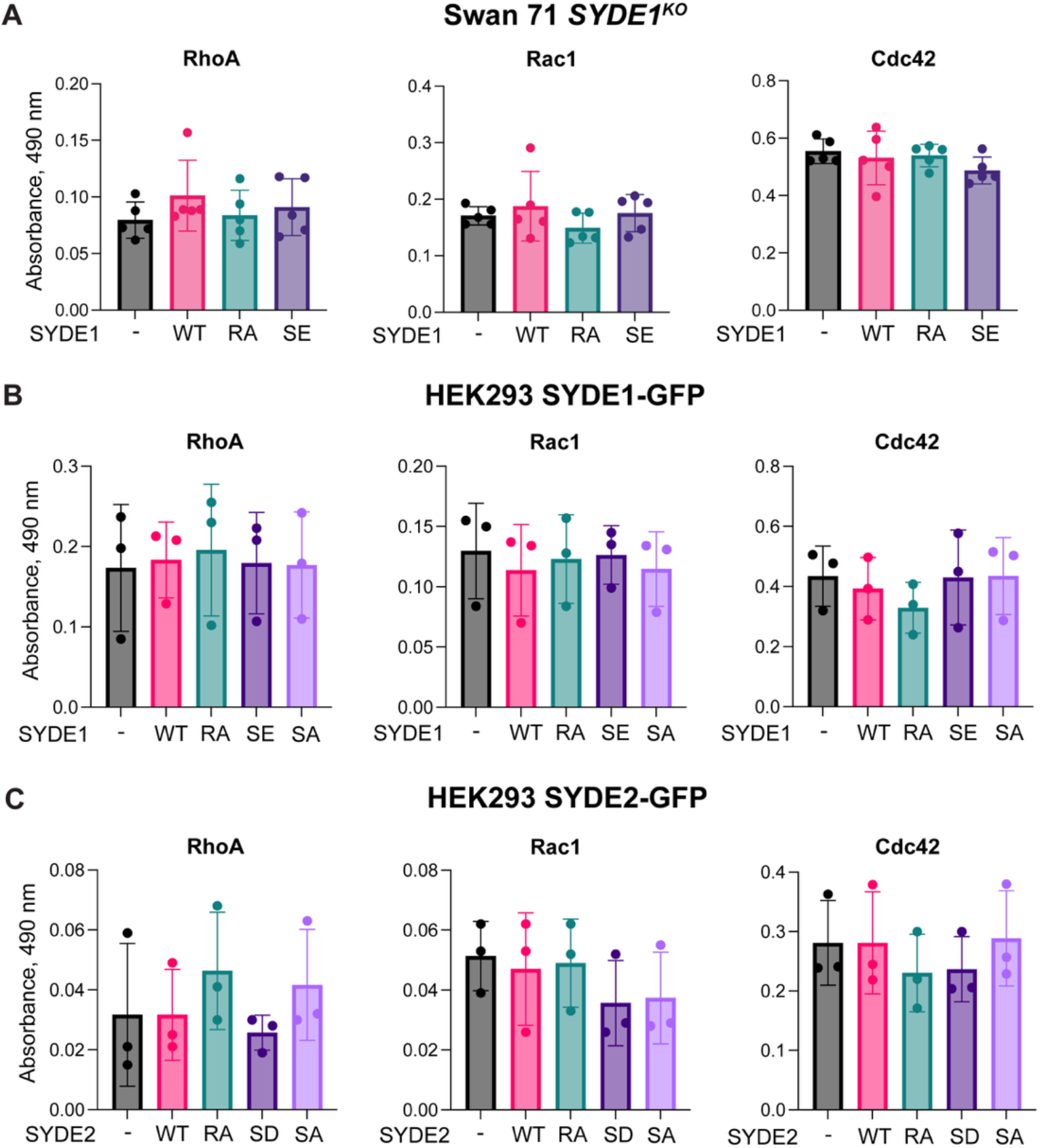
SYDE1 and SYDE2 do not significantly affect bulk GTP-loading of RhoA, Rac1, or Cdc42 in cells. (**A**) G-LISA effector binding assays for RhoA, Rac1, and Cdc42 in lysates from SYDE1 KO Swan 71 cells expressing the following constructs: GFP (−), SYDE1-GFP WT (WT), SYDE1-GFP R369A (RA), SYDE1-GFP S578E (SE). Data shown as mean ± SD from independent replicates (n = 5). All pairwise comparisons were insignificant (P >0.05) by ordinary one-way ANOVA with Tukey’s multiple comparisons test. (**B**) G-LISA effector interaction assays for RhoA, Rac1, and Cdc42 in lysates from HEK293 cells expressing GFP, SYDE1-GFP WT, RA, SE, and S578A (SA). Data shown as mean ± SD from independent replicates (n = 3). All pairwise comparisons were insignificant (P >0.05) by ordinary one-way ANOVA with Tukey’s multiple comparisons test. (**C**) G-LISA assays for RhoA, Rac1, and Cdc42 in lysates from HEK293 cells expressing GFP, SYDE2-GFP WT (WT), R707A (RA), S935D (SD), and S935A (SA). Data shown as mean ± SD from independent replicates (n = 3). All pairwise comparisons were insignificant (P >0.05) by ordinary one-way ANOVA with Tukey’s multiple comparisons test.

### SYDE1 and SYDE2 alter morphology in HEK293 cells

Rho GTPase signaling is established to regulate actin dynamics, and SYDE1 and SYDE2 have been implicated in cytoskeleton-related processes ^13,18^. Consistent with a possible effect on the actin cytoskeleton, we observed by fluorescence microscopy that HEK293 cells overexpressing GFP-tagged SYDE1 or SYDE2 displayed distinct morphology from control cells expressing GFP alone. Compared to expression of GFP, expression GFP-tagged SYDE1 or SYDE2 in HEK293 cells led to significantly increased cell area and decreased circularity (**Fig. 4A-C**). Quantitative analysis revealed that mutation of the catalytic Arg residue in SYDE1 significantly inhibited its ability to increase cell area, indicating that the GAP activity of SYDE1 promotes cell spreading (**Fig. 4B, left panel**). While there was a trend for cells expressing SYDE2 RA to have reduced spreading compared to SYDE2 WT, the effect was not statistically significant (**Fig. 4B, right panel**). Cells expressing SYDE1 RA and SYDE2 RA trended toward higher circularity compared to those expressing the corresponding WT forms, although this did not meet significance threshold in our analysis (**Fig. 4C**). These results indicate that SYDE1 and SYDE2 play a significant role in altering morphology when overexpressed in HEK293 cells, and that their catalytic GAP activity contributes partially to these functions.

**Fig. 4.**
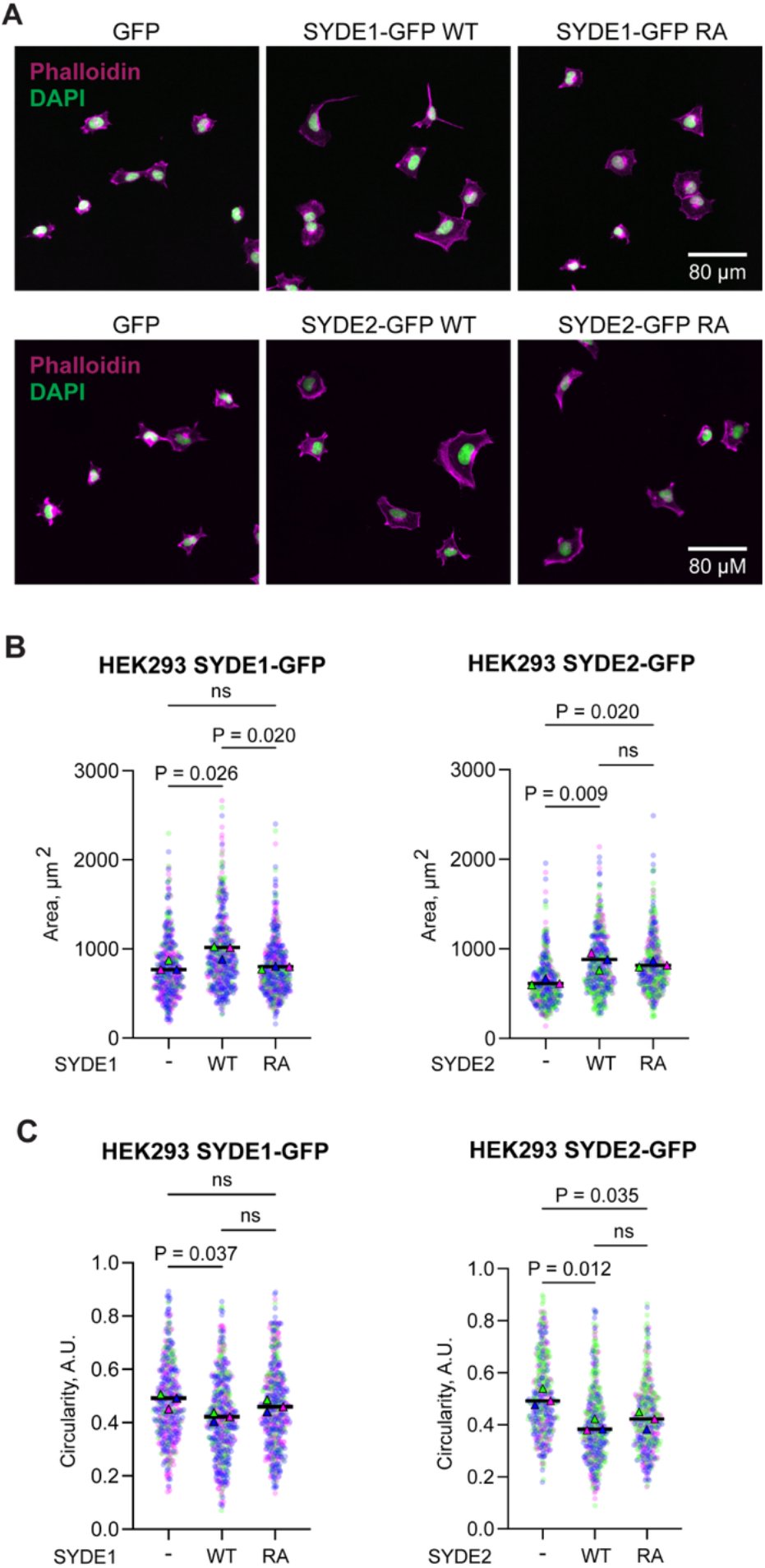
SYDE1 and SYDE2 alter cell area and circularity in HEK293 cells. (**A**) Representative widefield fluorescence images from n = 3 independent replicates of HEK293 cells expressing indicated constructs and stained with indicated markers. Merged images of phalloidin (magenta) and DAPI (green) staining are shown. Regions of overlapping fluorescence appear white. (**B**) Analysis of cell area from phalloidin staining segmentation quantified with ImageJ Fiji software. Data shown as mean values (triangles) overlaid on individual cell datapoints (circles) from n = 3 independent replicates of HEK293 cells expressing indicated constructs. Significance of differences in mean values was calculated using ordinary one-way ANOVA with Tukey’s multiple comparisons test. (**C**) Analysis of cell circularity from phalloidin staining segmentation quantified with ImageJ Fiji software. Data depicted and significance calculated as in panel (B). A.U. = Arbitrary Units.

### SYDE1 and SYDE2 increase the number of focal adhesions in HEK293 cells

Focal adhesions are key structures that mediate spreading and protrusion in cultured cells ^32^, and SYDE1 and SYDE2 reportedly localize to focal adhesion complexes ^13^. Based on the altered cell morphology we observed, we reasoned that SYDE1 and SYDE2 expression may affect focal adhesion formation or maintenance. Indeed, we found that HEK293 cells expressing GFP-tagged SYDE1 or SYDE2 had significantly increased numbers of focal adhesions per cell compared to those expressing GFP alone (**Fig. 5A, B**). Quantitative analysis revealed that unlike cells expressing WT SYDE1, cells expressing SYDE1 RA did not have significantly more focal adhesions than those expressing GFP, indicating that GAP activity of SYDE1 is important in this function. As we observed for its effect on cell morphology, there was a trend for SYDE2 RA to have a less severe phenotype compared to SYDE2 WT, but this effect did not meet our significance threshold. In contrast to focal adhesion number, we did not observe significant differences in focal adhesion area between cells expressing GFP-tagged SYDE1 or SYDE2 and those expressing GFP alone (**Fig. 5C**). Taken together, these results confirm that SYDE1 and SYDE2 play a role in promoting focal adhesions, which may underlie their capacity to influence cell shape.

**Fig. 5.**
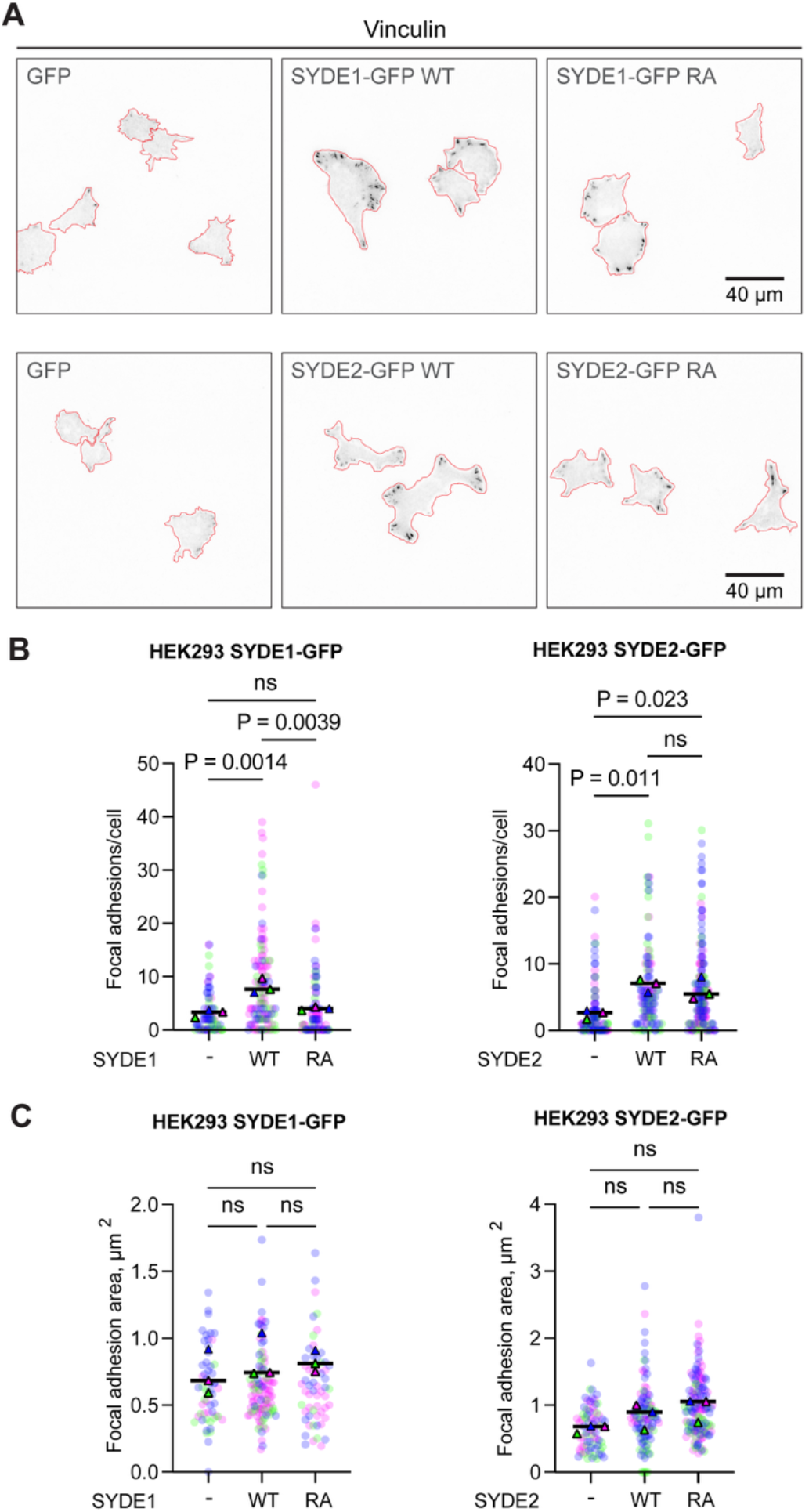
SYDE1 and SYDE2 increase number of focal adhesions in HEK293 cells. (**A**) Representative TIRF images of focal adhesions, visualized by vinculin immunostaining, from n=3 independent replicates of HEK293 cells expressing indicated constructs. For clarity, fluorescence is displayed using an inverted gray look-up table. Red outlines depict TIRF footprints. (**B**) Analysis of number of focal adhesions per cell from vinculin immunostaining quantified with ImageJ Fiji software. Data shown as mean values overlaid on individual cell data from n = 3 independent replicates of HEK293 cells expressing indicated constructs. Significance of differences in mean values was calculated using ordinary one-way ANOVA with Tukey’s multiple comparisons test. (**C**) Analysis of focal adhesion area per cell from vinculin immunostaining quantified with ImageJ Fiji software. Data depicted and significance calculated as in panel (B).

### SYDE1 and SYDE2 increase directional migration in HEK293 cells

Alterations in cell morphology and adhesion are intimately linked with migratory behavior ^33^. In order to assess whether the morphological and focal adhesion phenotypes in SYDE1- or SYDE2-expressing HEK293 cells correlate with migration, we tracked movement of individual cells over time by time-lapse imaging. We found that HEK293 cells expressing GFP-tagged SYDE1 or SYDE2 had significantly altered migration paths compared to cells expressing GFP alone. In general, HEK293 cells expressing GFP alone appear to have more confined and less persistently directional movement than cells expressing SYDE1 or SYDE2. Notably, cells expressing SYDE1 or SYDE2 showed decreased mean directional change rate compared to cells expressing GFP, which was significantly, though only partially, inhibited in the catalytic Arg mutants (**Fig. 6A**). Both SYDE1-expressing cells and SYDE2-expressing cells additionally had significantly increased linearity of forward progression compared to GFP-expressing cells (**Fig. 6B**), with SYDE1-expressing cells also showing significantly decreased total distance traveled and track mean speed (**Fig. 6C, D**). Possibly related to the partial effect on cell morphology, we did not observe significant differences between WT and the catalytic Arg mutant for any parameter other than mean directional change rate. Collectively, these results demonstrate that SYDE1 and SYDE2 can modulate migration behavior in HEK293 cells, which is associated with their functions in modulating cell morphology and focal adhesions.

**Fig. 6.**
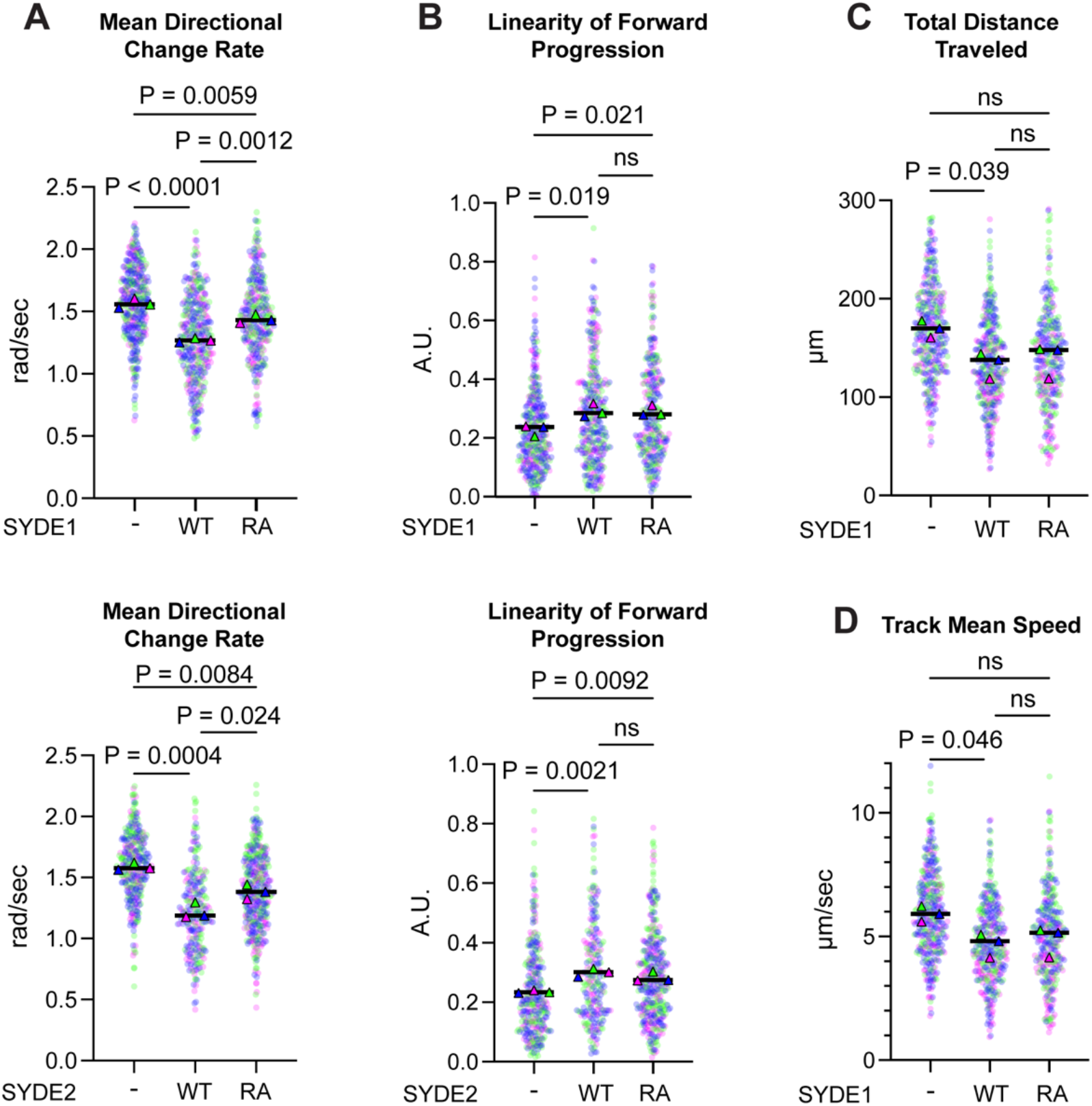
SYDE1 and SYDE2 alter migration in HEK293 cells. Analysis of (**A**) mean directional change rate, (**B**) linearity of forward progression, (**C**) total distance traveled, and (**D**) track mean speed quantified with ImageJ Fiji Trackmate plugin using GFP to threshold cell outlines. Data shown as mean values overlaid on individual cell data from n = 3 independent replicates of HEK293 cells expressing indicated constructs. Significance of changes in mean values was calculated using ordinary one-way ANOVA with Tukey’s multiple comparisons test. A.U. = Arbitrary Units.

## DISCUSSION

While many human RhoGAPs have been extensively characterized, SYDE1 and SYDE2 remain comparatively understudied. Our findings reveal biochemical and functional properties of SYDE1 and SYDE2 in vitro and in cells, and they implicate both proteins in the regulation of cell morphology, focal adhesions, and migration. We found that both SYDE1 and SYDE2 were robustly and selectively phosphorylated by JNK1 relative to other MAPKs at a specific Ser residue within each protein. This residue is conserved across nearly all vertebrate species ^34^, suggesting that SYDE1 and SYDE2 are regulated at some level by phosphorylation at this site. We further showed that mammalian SYDE1 and SYDE2 can act as GAPs on RhoA, Rac1, and Cdc42 in vitro, indicating that catalytic GAP activity could play a role in their biological functions. Consistent with this idea, the effects of SYDE1 and SYDE2 on cell morphology, focal adhesions, and migration were partially dependent on GAP activity, even though we could not detect a corresponding effect on total GTP loading of RhoA, Rac1, or Cdc42 in cells. Taken together, our study provides new insight into fundamental characteristics of SYDE1 and SYDE2 and identifies clear directions for further mechanistic investigation into their biological roles.

The functional significance of SYDE1 Ser578 and SYDE2 Ser935 phosphorylation by JNK remains an open question. MAP kinases are canonically activated downstream of Rho GTPases ^35–38^, raising the possibility that JNK phosphorylation of SYDE1 and SYDE2 participates in feedback regulation of Rho signaling. Phosphorylation is an established mechanism for regulating other GAP proteins, often modulating GAP activity indirectly by changing subcellular localization or intermolecular interactions ^4,39,40^. Aside from Rho GTPases, relatively little is known about SYDE1 and SYDE2 interacting proteins, and whether such interactions are regulated. Direct binding partners have been reported in neuronal contexts in *C. elegans*, *Drosophila*, and mice ^10,16,17^. An intramolecular interaction between the N-terminal tail/C2 domain of mammalian SYDE1 and its RhoGAP domain has been proposed to autoinhibit GAP activity ^12^. For human SYDE1 and SYDE2, high-throughput, low-throughput, and computational approaches have identified candidate binding partners ^13,34,41,42^, however phospho-modulation of these intermolecular interactions remains unexplored.

Direct phospho-regulation of GAP catalytic activity in vitro has been reported in only a limited number of cases ^23–28^. Our results examining GAP activity in vitro show that the phosphomimetic substitution at SYDE1 Ser578 slightly increases GAP activity toward RhoA. However, as phosphomimetic mutations are an imperfect substitute for true phosphorylation, this result does not in itself rule out a larger role for Ser578 phosphorylation in regulation of GAP activity. This experiment also used a construct of SYDE1 containing truncated N- and C-terminal unstructured regions (172-587), limiting potential regulatory mechanisms related to autoregulation or homodimerization. More conclusively, we found that JNK phosphorylation of full length SYDE2 does not directly alter its catalytic GAP activity toward RhoA, Rac1, and Cdc42 in vitro. However, these experiments do not rule out potential effects of JNK phosphorylation on GAP activity toward other Rho GTPases, nor do they rule out indirect effects on SYDE2 function through changes in subcellular localization or interaction with other proteins.

Perhaps surprisingly given their demonstrated GAP activity in vitro, we found that neither SYDE1 nor SYDE2 contributed significantly to global GTP-loading of RhoA, Rac1, or Cdc42 in Swan 71 trophoblasts or HEK293 cells. However, these negative results do not necessarily indicate that SYDE1 and SYDE2 GAP activity is functionally unimportant in cells. Because the assays we employed measure bulk GTP-loading across a population of cells, they cannot resolve spatially-restricted pools of Rho GTPase activation or activation confined to a sub-population of cells. A related issue is that the relatively high experimental variability associated with this technique would only allow for detection of large changes in overall GTP loading. Moreover, RhoGAPs can display activity toward Rho GTPases in vitro beyond their authentic targets in cells ^6^, and it therefore may be that RhoA, Rac1, and Cdc42 are not the primary cellular targets for SYDE1 and SYDE2. SYDE1 can reportedly bind to constitutively-active forms of RhoQ and RhoJ ^14^, though whether these are true cellular targets remains to be seen. Additionally, RhoC and the atypical Rho GTPase Rnd3/RhoE have been implicated in migration of HTR-8/SVneo trophoblast cells ^43,44^. Collectively, these studies raise the possibility that SYDE1 and SYDE2 target Rho family members outside of RhoA, Rac1, and Cdc42. Taken together with our findings, these studies indicate that better defining the cognate Rho GTPases for SYDE1 and SYDE2 in cells is an important future direction.

We revealed functions of SYDE1 and SYDE2 in regulating cell morphology, focal adhesions, and migration, supporting previous studies suggesting them to have roles related to the actin cytoskeleton ^13,18^. It is well established that Rho GTPase signaling governs the dynamics of actin filament polymerization and organization that underlie fundamental cellular functions ^1,2^. Classical literature attributes stress fiber formation, actomyosin contractility, and focal adhesion maturation to RhoA ^45^, lamellipodia formation and membrane ruffling to Rac1 ^46^, filopodia formation and cell polarity to Cdc42 ^47,48^, and transient adhesion complex formation to Rac1 and Cdc42 ^48^. This framework is now understood as an oversimplification of a complex signaling network in which cellular functions rely on sequential activation, deactivation, or co-activation of multiple Rho GTPase family members, and precise spatiotemporal inactivation by RhoGAPs is essential for this regulation ^49–53^. Because SYDE1 and SYDE2 can promote RhoA, Rac1, and Cdc42 GTP hydrolysis in vitro, and because they contribute to at least partially GAP-dependent cellular phenotypes, it is possible that SYDE1 and SYDE2 contribute to spatiotemporal inactivation of distinct pools of these Rho GTPases. However, the exact mechanisms by which this might occur and the potential contribution of non-catalytic functions of SYDE1 and SYDE2 to these phenotypes remain to be defined. In summary, our study establishes fundamental biochemical and functional characteristics of SYDE1 and SYDE2 which provide a foundation for further investigation into their biological functions and regulation.

## MATERIALS AND METHODS

### Plasmids, cloning, and mutagenesis

The pV1900-SYDE1-3xFlag and pcDNA3-flag-SYDE2 mammalian expression vectors for C-terminal 3x flag-tagged full-length human SYDE1 (UniProt: Q6ZW31-2, isoform 2) and N-terminal flag-tagged full-length mouse SYDE2 (UniProt: A0A0G2JEY9) were previously generated ^20^. A codon-optimized pET28a-SYDE1 bacterial expression vector for N-terminal 6xHis-tagged human SYDE1 (residues 172-587) was custom produced by Biomatik. The pET28a-SYDE2 bacterial expression vector for N-terminal 6x His-tagged mouse SYDE2 (residues 507-944) was made by subcloning the coding sequence from pcDNA3-flag-SYDE2 into the BamHI and NotI restriction sites. The pGEX-SYDE2-His6 bacterial expression vector for dual-tagged (N-terminal GST and C-terminal 6xHis) SYDE2 (residues 507-944) was generated by Gibson assembly. The vectors pLX304-SYDE1-GFP and pLX304-SYDE2-GFP for lentiviral expression of C-terminally GFP-tagged SYDE1 and SYDE2 respectively were generated by Gateway cloning as follows. A Gateway pDONR 221 entry vector was made by BP recombination of attB1/B2-flanked SYDE1-GFP or SYDE2-GFP sequences (generated by overlap extension PCR of the SYDE1 sequence from pV1900-SYDE1-3xFlag and the SYDE2 sequence from pcDNA3-flag-SYDE2 with a GFP coding sequence from pEGFP-N1), which was then recombined by LR reaction into a pLX304 Gateway destination vector. A pLX304-GFP control vector was generated similarly. All point mutants were generated by QuikChange (Agilent) site-directed mutagenesis following manufacturer protocols. All constructs were confirmed by Sanger sequencing through the entire open reading frame. pGEX-6p1-RhoA, pET28a-Cdc42, and pET28a-Rac1 bacterial expression vectors for human RhoA (residues 1-181), Cdc42 (residues 2-191), and Rac1 (residues 2-177), respectively, were gifts from the laboratory of Titus Boggon (Yale University).

### Protein Expression and Purification

For purification of untagged RhoA, BL21-Gold(DE3) *E. coli* cells (Agilent #230132) were transformed with pGEX-6p1-RhoA and grown in 1 L of Terrific Broth (TB) in a shaking incubator at 37 °C to an approximate OD_600_ of 0.8. Cultures were then cooled to 16°C, expression was induced with 0.5 mM isopropyl β-D-thiogalactopyranoside (IPTG), and shaking at 16°C incubator continued for 16 hr. Cells were pelleted at 2,000 *x g* for 15 min at 4°C, snap frozen on dry ice with 95% ethanol, thawed, and resuspended in 20 mL ice cold buffer containing 20 mM Tris, pH 7.5, 140 mM NaCl, 1 mM dithiothreitol (DTT), and cOmplete™ protease inhibitor tablet (Roche #11697498001). Resuspended cells were lysed by addition of 200 μg/mL lysozyme, 0.4% Igepal CA-630, 30 U/mL DNAse I, and 13 mM MgCl_2_ followed by rotation for 30 minutes at 4°C and sonication. Lysate was spun 40 min at 14,000 x *g* at 4°C, and supernatant was incubated with 0.25 mL pre-equilibrated glutathione Sepharose 4B affinity resin (Cytiva) for 1 hr at 4°C with rotation. Beads were pelleted for 2 min at 1,000 *x g* at 4°C, and supernatant was decanted before washing beads twice (5 minutes each with rotation at 4°C) with ice cold 10 mL wash buffer (20 mM Tris, pH 7.5, 140 mM NaCl, and 1 mM DTT). Beads were resuspended in 2 mL wash buffer and transferred to a gravity flow column, and protein was eluted by overnight incubation with GST-tagged HRV 3C protease with rotation at 4°C. Untagged RhoA was collected as flow-through the following day and further eluted at 4°C in 500 μL fractions with wash buffer. Fractions containing protein were combined and subjected to size exclusion chromatography on a Superdex 200 10/300 GL column (Cytiva) in 20 mM Tris, pH 8.0, 150 mM NaCl, and 1 mM DTT. Fractions containing eluted protein were combined and spin-concentrated in an Amicon Ultra-4 centrifugal filter (Millipore #801024) with a 10 kDa molecular weight cutoff and stored at −80°C until use. Protein purity and concentration were assessed by SDS-PAGE alongside BSA standards with Coomassie staining.

For purification of Cdc42, BL21-Gold(DE3) *E. coli* were transformed with pET28a-Cdc42 and grown as described for RhoA above, and cell pellets were resuspended in the same buffer containing 10 mM imidazole. Resuspended cells were lysed, processed, and spun in the same way, and supernatant was incubated with 0.25 mL of pre-equilibrated ProBond™ Nickel Chelating Resin (Invitrogen #R80101) for 1 hr at 4°C with rotation. Beads were centrifuged for 2 minutes at 1,000 *x g* at 4°C and supernatant was decanted before resuspending in 5 mL of ice cold PBS with 0.5% Igepal CA-630 and transferring to a 10 mL gravity flow column at 4°C. Suspension was drained and washed once with 4 mL of the same buffer, then washed once with 4 mL of ice cold wash buffer containing 20 mM Tris pH 7.5, 150 mM NaCl, 10 mM imidazole, 0.01% Igepal CA-630, and 1 mM DTT. Protein was eluted in 0.5 mL fractions of ice cold elution buffer containing 20 mM Tris pH 7.5, 150 mM NaCl, 250 mM imidazole, and 1 mM DTT. Fractions containing protein were combined and fractionated by size exclusion chromatography as described above for RhoA.

For purification of Rac1, Rosetta™ 2(DE3) E. coli (EMD Millipore #71400-4) were transformed with pET28a-Rac1 and grown in 3 L of TB in a shaking incubator at 37°C to an OD_600_ of 0.6, cooled to 30°C, induced with 0.5 mM IPTG, and shaking at 30°C continued for 5 hr. Cells were processed as for RhoA purification above, and resuspended in 30 mL ice cold buffer containing 20 mM Tris pH 7.5, 140 mM NaCl, 3 mM β-mercaptoethanol, cOmplete™ protease inhibitor tablet, and 10 mM imidazole. Resuspended cells were lysed, processed, and spun as described above for RhoA, and supernatant was incubated with 0.75 mL of pre-equilibrated Talon® immobilized metal affinity resin (Takara Bio, #635502) for 1 hr at 4°C with rotation. Beads were pelleted, resuspended, transferred to a gravity flow column, drained, and washed once with 4 mL of ice cold PBS with 0.5% Igepal CA-630, and once with 4 mL wash buffer containing 20 mM Tris pH 7.5, 150 mM NaCl, 10 mM imidazole, 0.01% Igepal CA-630, and 3 mM β-mercaptoethanol. Protein was eluted in 1.5 mL fractions with a step-wise imidazole gradient of buffer containing 20 mM Tris pH 7.5, 150 mM NaCl, 3 mM β-mercaptoethanol, and imidazole at concentrations of 25 mM, 50 mM, 75 mM, 100 mM, 125 mM, 150 mM, 175 mM, 200 mM, 225 mM, 250 mM, or 300 mM. Fractions containing protein as assessed by SDS-PAGE and Coomassie staining were combined and exchanged into final buffer containing 20 mM Tris pH 8.0, 150 mM NaCl, and 1 mM DTT on PD midiTrap G-25 gravity de-salting columns (Cytiva #28918008). Sample was spin-concentrated, protein quantified as described above, and stored at −80°C until use.

WT and mutant variants of 6xHis-SYDE1 (residues 172-587) and 6xHis-SYDE2 (residues 507-944) were expressed in BL21-Gold(DE3) E. coli and grown as described above for Rac1, then cooled and induced as described above for RhoA. Protein was then purified on Talon resin as described above for Rac1, importantly without rotation for the resin-binding step.

Doubly tagged WT and mutant GST-SYDE2-6xHis (residues 507-944) was sequentially purified on glutathione resin and immobilized metal affinity chromatography as follows. Protein was expressed in Rosetta™ 2(DE3) *E. coli*, which was grown as described above for Rac1, then induced with 0.5 mM IPTG at 37°C shaking for 3 hours, lysed, and bound on glutathione Sepharose 4B beads as described for RhoA above. Beads were pelleted, washed twice in batch with 5 mL ice cold PBS with 0.5% Igepal CA-630, and once with 5 mL of ice cold wash buffer containing 50 mM Tris pH 8.0, 50 mM NaCl, 1 mM DTT, 0.01% Igepal CA-630, and 10% glycerol. Beads were resuspended in 1 mL wash buffer and transferred to an Eppendorf tube, pelleted, and protein was eluted in batch in two fractions of 0.5 mL wash buffer with 20 mM reduced glutathione (pH 7.0). Fractions were combined and exchanged into the resuspension buffer used for purification of Rac1 above. Samples were purified with a stepwise imidazole gradient, de-salted, and concentrated as above, and stored at −80°C until use. Flag-SYDE2 WT and RA were purified from HEK293T cells (ATCC #CRL-3216) transfected with pcDNA3-flag-SYDE2 using polyethyleneimine (PEI) and purified as previously described ^54^. Cells were treated for 1 hour before harvesting with either 0.1% DMSO vehicle control, anisomycin (10 μg/mL), or JNK-IN-8 (5 μM).

### Cell culture

HEK293 and HEK293T cells were cultured in DMEM (Gibco #11965-092) supplemented with 10% fetal bovine serum (Gibco #A52567-01). Swan 71 cells were a gift from the laboratory of Gil Mor (Yale University, New Haven, CT, USA) and cultured in DMEM supplemented with 10% fetal bovine serum, 10mM HEPES (Gibco #15630-080), 1 mM sodium pyruvate (Gibco #11360-070), and 1x MEM NEAA (Gibco #11140-050).

### Cell lysis, immunoblotting, and antibodies

To process samples for immunoblotting, cells were washed twice with cold PBS and lysed on ice for 10 minutes with buffer containing 20 mM Tris pH 7.5, 150 mM NaCl, 1% Triton X-100, 1 mM β-glycerophosphate, 2.5 mM sodium pyrophosphate, 1 mM Na_3_VO_4_, and cOmplete™ protease inhibitor tablet. Cell lysates were clarified by spinning at 10,000 *x g* for 10 minutes in a microcentrifuge at 4°C. Protein concentration of clarified lysates was determined by BCA protein assay and 4x SDS-PAGE loading buffer was added to samples. Samples were heat denatured, subjected to SDS-PAGE and transferred to polyvinyl difluoride (PVDF) membranes (Immobilon®-FL 0.45 μm pore size, Millipore #IPFL00010). Membranes were blocked for 1 hour at room temperature in Tris buffered saline with 0.1% Tween20 (TBS-T) with 5% nonfat dry milk and incubated overnight at 4°C in primary antibodies diluted according to manufacturer’s instructions. The following day, membranes were washed 3 × 10 minutes with TBS-T before incubation with fluorophore-conjugated secondary antibodies diluted 1:10,000 in blocking buffer for 1 hour at room temperature. Membranes were then washed 3 × 10 minutes with TBS-T, analyzed on an Odyssey CLx LI-COR imager, and quantified with Image Studio Lite software.

The following primary antibodies were used for immunoblotting: SYDE1 (Atlas Antibodies #HPA013328), GFP (Rockland #600-101-215), Flag (Sigma-Aldrich #F3165), vinculin (Sigma-Aldrich #V9131), β-actin (Sigma-Aldrich #A5441), RhoA (Cell Signaling Technology #2117), Rac1 (Cell Signaling Technology #2465), Cdc42 (Cell Signaling Technology #2466). The rabbit polyclonal anti-SYDE1 phosphospecific antibody was raised against a phosphopeptide spanning the phosphorylated Ser578 residue (RGRGGPE**<u>S</u>**PPSNRYA) by YenZym Antibodies. The phosphospecific antibody was affinity purified through iterative rounds of subtraction and affinity purification as previously described ^55^ with the unphosphorylated and phosphorylated peptide, respectively. Fluorophore-conjugated secondary antibodies used for immunoblotting were: IRDye® 800CW donkey anti-rabbit IgG (LI-COR #926-32213), IRDye® 680RD donkey anti-goat IgG (LI-COR #926-68074), IRDye® 800CW donkey anti-mouse IgG (LI-COR #926-32212), Alexa Fluor™ 680 goat anti-rabbit IgG (Invitrogen #A21109).

### Analysis of SYDE1 phosphorylation in cultured cells by mass spectrometry

HEK293T cells were transiently transfected with pV1900-SYDE1-3xFlag plasmid using PEI, treated, and processed, and subjected to MS analysis as previously described ^20^ at the Keck Mass Spectrometry & Proteomics Resource (Yale University).

### Protein kinase assays

Relative activities of JNK1 (Sino Biological #M33-10G), p38α (Reaction Biology #0443-0000-3), ERK5 (CarnaBio #04-146), and ERK2 (purified previously from *E. coli* ^56^) were assessed in fluorogenic kinase assays using a sulfonamido-oxine (SOX)-containing MAPK substrate peptide (AssayQuant, AQT0376). Assays were performed in biological duplicate in 96-well half area white flat-bottomed microplates (Corning #3693). Peptide substrate was diluted to 5 μM in reaction buffer containing 54 mM HEPES pH 7.5, 1 mM ATP, 1 mM DTT, 0.012% Brij-35, 1% glycerol, 10 mM MgCl_2_, 0.2 mg/mL BSA, and 0.55 mM EGTA and pre-incubated 5 min at 30°C. Reactions were initiated by addition of kinase in 5x kinase dilution buffer (20 mM HEPES pH 7.5, 1 mM DTT, 0.01% Brij-35, 5% glycerol, 1 mg/mL BSA, and 0.1 mM EGTA) with final kinase concentrations of JNK1 = 5 nM, p38 = 5 nM, ERK5 = 5 nM, and ERK2 = 0.5 nM. Immediately after addition of kinase, fluorescence (excitation 360 nm, emission 485 nm) was read every 90 seconds over 120 minutes on a Synergy H1 Microplate Reader with Gen5 software (v3.11.19)(BioTek) at 30°C. Reaction rates were calculated from the linear portion of the reaction progress curve to define units of kinase activity. Equal units of activity of each kinase were added to subsequent protein kinase assays with SYDE1 and SYDE2.

WT and SA mutant forms of SYDE1 (residues 172-587, 350 nM) were incubated with kinases at concentrations corresponding to equivalent units of activity (JNK1, 50 nM; p38α, 3.3 nM; ERK5, 35.7 nM; ERK2, 1.5 nM) in reaction buffer containing 20 mM Tris pH 7.5, 150 mM NaCl, and 1 mM DTT. Reactions were initiated by addition of MgCl_2_ and ATP to a final concentrations of 1 mM and 25 μM, respectively, and incubated at 30°C for 15 min. Reactions were quenched by addition of 4x SDS-PAGE loading buffer, subjected to SDS-PAGE, and immunoblotted with SYDE1 pSer578 antibody. Reactions with GST-SYDE2(507-944)-6xHis (SYDE2, 140 nM; JNK1, 30 nM; p3, 2 nM; ERK5, 21 nM; ERK2, 0.91 nM) were conducted in reaction buffer containing 50 mM HEPES pH 7.4, 100 mM NaCl, 0.1 mM Na_3_VO_4_, 0.012% Brij-35, 10 mM MgCl_2_, and 1 mM DTT. Reactions were initiated by addition of [γ-^32^P]ATP (Revvity) to 20 μM at 0.1 μCi/μL and incubated 30 min at 30°C. Reactions were quenched by addition of 4x SDS-PAGE loading buffer separated by SDS-PAGE (7.5% acrylamide), stained with Coomassie, destained, and exposed on a phosphor screen. Autoradiographs were analyzed on an Amersham Typhoon phosphorimager and quantified with ImageJ Fiji software ^57^.

### GTPase activity assays

Multiple turnover endpoint RhoA, Rac1, and Cdc42 assays were conducted using BIOMOL Green (Enzo BML-AK111) malachite green reagent in transparent, flat-bottomed 96-well microplates. Each reaction mixture contained 5 μM of GTPase, 500 nM of 6xHis-SYDE1, 25 nM of 6xHis-SYDE2, or 25nM of Flag-SYDE2, in reaction buffer (20 mM Tris pH 8.0, 250 mM NaCl, 1 mM MgCl_2_, and 2 mM EDTA), and 76 μL of this reaction mixture was added to each well. To initiate reactions, 4 μL of 1 mM GTP was added to each well and incubated at room temperature. Reactions were quenched with 100 μL of BIOMOL Green reagent according to manufacturer protocol. Absorbance at 620 nm was read in a Synergy H1 Microplate Reader with Gen5 software v3.11.19 (BioTek). Quantity of free phosphate (P_i_) released was quantified according to manufacturer protocol. Data were fit to linear rates of µM P_i_ released per min with Prism v10.1.1 (GraphPad Software, MA). Statistical significance was calculated using ordinary one-way ANOVA with Tukey’s multiple comparisons test.

### Generation of *SYDE1* KO Swan 71 cells

Single guide RNA (sgRNA) sequences targeting *SYDE1* exon 3 (TGTCCTACCTCGGAAGACCG) or exon 5 (CAGTGATGACATTGATATCG) were cloned into the Cas9 transient expression plasmid pSpCas9(BB)-2A-GFP (PX458, Addgene #48138) according to the cloning protocol established by the laboratory of Feng Zhang (The Broad Institute, Cambridge, MA, USA). Swan 71 cells were transfected with plasmids including *SYDE1*-targeting sgRNA or empty vector with PEI, and 48 hours post-transfection single GFP-positive cells were sorted into 96-well tissue culture plates using a BD FACSAria instrument at the Yale Flow Cytometry Facility. Clones were expanded, and *SYDE1* KO was confirmed by immunoblotting with SYDE1 antibody. Single clonal control and confirmed *SYDE1* KO cell lines were selected for subsequent reconstitution experiments.

### Generation of SYDE1-GFP- and SYDE2-GFP-expressing stable cell lines

Swan 71 and HEK293 cells stably expressing GFP alone, or WT or mutant forms of SYDE1-GFP or SYDE2-GFP were generated by transduction with lentivirus generated from pLX304-GFP, pLX304-SYDE1-GFP, and pLX304-SYDE2-GFP, respectively. Lentiviral plasmids were packaged by PEI co-transfection in HEK293T cells with dR8.91 packaging plasmid and VsV-G envelope protein at a pLX304:dR8.91:VsV-G ratio of 10:10:1. After 48 hours, virus-containing medium was harvested, filtered, and stored at −80°C. Target cells were transduced in the presence of 8μg/mL polybrene for 24 hr followed by selection for >10 days in media containing 10 μg/mL blasticidin S (Gibco #A1113903). GFP-positive cells were bulk sorted using a BD FACSAria instrument at the Yale Flow Cytometry Facility. Protein expression was confirmed by immunoblotting with GFP and SYDE1 antibodies.

### Cell-based GTP loading assays

GTP-bound GTPase levels in cell lysates were measured using G-LISA assay kits for RhoA (Cytoskeleton, Inc. #BK124), Rac1 (Cytoskeleton, Inc. #BK128), and Cdc42 (Cytoskeleton, Inc. #BK127). Swan 71 or HEK293 cells were seeded at a density of 400,000 or 700,000 cells respectively per 10 cm dish. HEK293 cells were harvested 24 hr after seeding, while Swan 71 cells were exchanged into serum-free media 24 hr after seeding and harvested the following day. Lysates for G-LISA assays were prepared according to manufacturer’s protocols. Lysates were quantified with Precision Red Advanced Protein Assay reagent according to manufacturer’s protocols and protein concentrations were equalized before subjecting to G-LISA assay. Absorbance was read at 490 nm with a Synergy H1 Microplate Reader with Gen5 software, and data were graphed in Prism v10.1.1. Statistical significance was calculated using ordinary one-way ANOVA with Tukey’s multiple comparisons test.

### Immunofluorescence microscopy

HEK293 cells stably expressing indicated constructs were plated on fibronectin-coated (1 μg/cm^2^, Sigma Aldrich #F1141) 35 mm glass-bottom dishes (Cellvis #D35-20-1.5-N) 12 hours prior to 15 min room temperature fixation with 4% paraformaldehyde (Electron Microscopy Sciences #15710) in 1x DPBS (Gibco #14190-144). Cells were washed 3 x 5 minutes with 1x DPBS before blocking 1 hr with immunofluorescence blocking buffer (Cell Signaling Technology #12411) at room temperature. Vinculin (Sigma-Aldrich #V9131) primary antibody diluted 1:1000 in immunofluorescence antibody dilution buffer (Cell Signaling Technology #12378) was added to cells and incubated overnight at 4°C. The following day, cells were washed 3 x 5 min with 1x DPBS before incubation for 1 hr at room temperature with Alexa Fluor™ 568 goat anti-mouse IgG (Invitrogen #A11031) and Alexa Fluor™ 647 phalloidin (Invitrogen #A22287) diluted 1:1000 and 1:200, respectively, in immunofluorescence antibody dilution buffer. Lastly, cells were washed 3 x 5 minutes with 1x DPBS, with 3.3 μg/mL DAPI (Thermo Fisher Scientific #62248) included in the second-to-last wash. Samples were stored in 1x DPBS overnight before imaging on the Dragonfly 620-SR System (Andor Technology) equipped with a Nikon ECLIPSE Ti2 inverted microscope (Nikon Instruments) with Perfect Focus System. For cell morphology and migration experiments, cells were imaged by widefield microscopy using a 20x objective (air). For focal adhesion studies, cells were imaged using objective-type total internal reflection fluorescence (TIRF) microscopy with a 60x objective (NA 1.49, oil) and Borealis illumination (Andor Technology) for uniform evanescent field. Images were captured on a scientific complementary metal oxide semiconductor (sCMOS) camera (Sona 4.2B-6, Andor Technology). Fields were acquired as 4×4 tiled images and stitched with Fusion software (version 2.6.1, Andor Technology) to make finalized raw image files.

### Live cell microscopy

Time-lapse images of HEK293 cells stably expressing indicated constructs were plated in phenol red-free DMEM supplemented with glucose, L-glutamine, and HEPES (Gibco #21063-029) and additionally supplemented with 10% fetal bovine serum, 1 mM sodium pyruvate, and 1x MEM NEAA on fibronectin-coated (1 μg/cm^2^) glass-bottom 8-well slides (Ibidi μ-Slide #80807) 12 hr prior to imaging. Tiled 4×4 images were acquired at 20x wide field magnification every 5 minutes for 2.5 hours on a Dragonfly spinning-disk confocal microscope with a chamber around the microscope stage maintained at 37°C and 5% CO_2_. Images were stitched with Imaris software to make finalized raw movie files.

### Cell morphology, focal adhesion, and migration analysis

Cell morphology was analyzed with ImageJ Fiji software. In brief, raw microscopy images were qualitatively matched for cell density and blinded, and cell outlines were created by thresholding and segmenting the image using phalloidin staining signal to create a binary mask following cell perimeters. Segmented images were then manually inspected to exclude outlines corresponding to clumped cells, debris, and artifacts. At least 91-206 individual cell measurements were obtained per cell line across all three independent replicates. The mean value of individual cell measurements from each cell line was used to calculate statistical significance using ordinary one-way ANOVA with Tukey’s multiple comparisons test on Prism.

Focal adhesions were analyzed with ImageJ Fiji software as previously described ^58^. In brief, raw microscopy images were first converted from 16-bit to 8-bit and background subtracted using the rolling ball radius set to 50 pixels with the sliding paraboloid option. Contrast was then enhanced using the plugin CLAHE (Contrast Limited Adaptive Histogram Equalization) with block size set at 19, histogram bins set at 256 and maximum slope set at 6. Background was then further minimized by applying the “mathematical exponential (Exp)” function in ImageJ. Finally, processed images were thresholded and focal adhesions were quantified using the “analyze particle” function with the minimum area set to 0.2 mm^2^.

Cell migration tracks were analyzed with the Trackmate plugin for ImageJ Fiji software ^59,60^. In brief, raw microscopy movies were thresholded on Trackmate to create outlines around cell perimeters, which was used to draw tracks corresponding to the migration of individual cells. Cells and tracks were then manually inspected to exclude tracks corresponding to clumped cells, debris, artifacts, and other factors (cell division, cell collision, apoptosis) that resulted in a compromised track or an incomplete track for the duration of the movie. At least 338-387 individual cell track measurements were obtained per cell line across all three independent replicates. The mean value of individual cell measurements from each cell line was used to calculate statistical significance using ordinary one-way ANOVA with Tukey’s multiple comparisons test on Prism.

## ACKNOWLEDGMENTS

We thank Titus Boggon, David Calderwood, Vikki Abrahams, Anthony Koleske, and members of their laboratories (Yale University), as well as members of the Turk laboratory for advice and suggestions regarding this work. We thank TuKiet Lam for LC-MS/MS analysis and Florine Collin and Jean Kanyo for assistance with proteomics sample preparation and data collection (Yale Keck Mass Spectrometry and Proteomics Resource). We thank Yale Flow Cytometry for their assistance with cell sorting. The Flow Cytometry Core is supported in part by an National Cancer Institute Cancer Center Support Grant NIH P30 CA016359. This work was supported by National Institutes of Health grants R35 GM153265 to B.E.T. C.S. was supported by National Institutes of Health fellowship F31 HD111285 and training grant T32 GM007324.

## REFERENCES

1. Etienne-Manneville, S. & Hall, A. Rho GTPases in cell biology. Nature 420, 629–635 (2002).

2. Jaffe, A. B. & Hall, A. RHO GTPASES: Biochemistry and Biology. Annu. Rev. Cell Dev. Biol. 21, 247–269 (2005).

3. Hall, A. Rho GTPases and the Actin Cytoskeleton. Science 279, 509–514 (1998).

4. Hodge, R. G. & Ridley, A. J. Regulating Rho GTPases and their regulators. Nat. Rev. Mol. Cell Biol. 17, 496–510 (2016).

5. Schaefer, A., Reinhard, N. R. & Hordijk, P. L. Toward understanding RhoGTPase specificity: structure, function and local activation. Small GTPases 5, e968004 (2014).

6. Mosaddeghzadeh, N. & Ahmadian, M. R. The RHO Family GTPases: Mechanisms of Regulation and Signaling. Cells 10, 1831 (2021).

7. Dahmene, M., Quirion, L. & Laurin, M. High Throughput strategies Aimed at Closing the GAP in Our Knowledge of Rho GTPase Signaling. Cells 9, 1430 (2020).

8. Amin, E. et al. Deciphering the Molecular and Functional Basis of RHOGAP Family Proteins. J. Biol. Chem. 291, 20353–20371 (2016).

9. Hallam, S. J., Goncharov, A., McEwen, J., Baran, R. & Jin, Y. SYD-1, a presynaptic protein with PDZ, C2 and rhoGAP-like domains, specifies axon identity in C. elegans. Nat. Neurosci. 5, 1137–1146 (2002).

10. Xu, Y., Taru, H., Jin, Y. & Quinn, C. C. SYD-1C, UNC-40 (DCC) and SAX-3 (Robo) Function Interdependently to Promote Axon Guidance by Regulating the MIG-2 GTPase. PLOS Genet. 11, e1005185 (2015).

11. Spinner, M. A., Walla, D. A. & Herman, T. G. *Drosophila* Syd-1 Has RhoGAP Activity That Is Required for Presynaptic Clustering of Bruchpilot/ELKS but Not Neurexin-1. Genetics 208, 705–716 (2018).

12. Wentzel, C. et al. mSYD1A, a Mammalian Synapse-Defective-1 Protein, Regulates Synaptogenic Signaling and Vesicle Docking. Neuron 78, 1012–1023 (2013).

13. Müller, P. M. et al. Systems analysis of RhoGEF and RhoGAP regulatory proteins reveals spatially organized RAC1 signalling from integrin adhesions. Nat. Cell Biol. 22, 498–511 (2020).

14. Bagci, H. et al. Mapping the proximity interaction network of the Rho-family GTPases reveals signalling pathways and regulatory mechanisms. Nat. Cell Biol. 22, 120–134 (2020).

15. Amado-Azevedo, J. et al. A CDC42-centered signaling unit is a dominant positive regulator of endothelial integrity. Sci. Rep. 7, 10132 (2017).

16. Owald, D. et al. A Syd-1 homologue regulates pre- and postsynaptic maturation in *Drosophila*. J. Cell Biol. 188, 565–579 (2010).

17. Owald, D. et al. Cooperation of Syd-1 with Neurexin synchronizes pre-with postsynaptic assembly. Nat. Neurosci. 15, 1219–1226 (2012).

18. Lo, H. et al. Association of dysfunctional synapse defective 1 (SYDE1) with restricted fetal growth – SYDE1 regulates placental cell migration and invasion. J. Pathol. 241, 324–336 (2017).

19. Jaju Bhattad, G., et al. Histone deacetylase 1 and 2 drive differentiation and fusion of progenitor cells in human placental trophoblasts. Cell Death Dis. 11, 311 (2020).

20. Shi, G. et al. Proteome-wide screening for mitogen-activated protein kinase docking motifs and interactors. Sci. Signal. 16, eabm5518 (2023).

21. Zhang, T. et al. Discovery of Potent and Selective Covalent Inhibitors of JNK. Chem. Biol. 19, 140–154 (2012).

22. Johnson, J. L. et al. An atlas of substrate specificities for the human serine/threonine kinome. Nature 613, 759–766 (2023).

23. Tripathi, B. K. et al. CDK5 is a major regulator of the tumor suppressor DLC1. J. Cell Biol. 207, 627–642 (2014).

24. Minoshima, Y. et al. Phosphorylation by Aurora B Converts MgcRacGAP to a RhoGAP during Cytokinesis. Dev. Cell 4, 549–560 (2003).

25. Bastos, R. N., Penate, X., Bates, M., Hammond, D. & Barr, F. A. CYK4 inhibits Rac1-dependent PAK1 and ARHGEF7 effector pathways during cytokinesis. J. Cell Biol. 198, 865–880 (2012).

26. Breznau, E. B., Semack, A. C., Higashi, T. & Miller, A. L. MgcRacGAP restricts active RhoA at the cytokinetic furrow and both RhoA and Rac1 at cell–cell junctions in epithelial cells. Mol. Biol. Cell 26, 2439–2455 (2015).

27. Walkup, W. G. et al. Phosphorylation of Synaptic GTPase-activating Protein (synGAP) by Ca2+/Calmodulin-dependent Protein Kinase II (CaMKII) and Cyclin-dependent Kinase 5 (CDK5) Alters the Ratio of Its GAP Activity toward Ras and Rap GTPases. J. Biol. Chem. 290, 4908–4927 (2015).

28. Walkup, W. G., Sweredoski, M. J., Graham, R. L., Hess, S. & Kennedy, M. B. Phosphorylation of synaptic GTPase-activating protein (synGAP) by polo-like kinase (Plk2) alters the ratio of its GAP activity toward HRas, Rap1 and Rap2 GTPases. Biochem. Biophys. Res. Commun. 503, 1599–1604 (2018).

29. Tripathi, B. K. et al. Receptor tyrosine kinase activation of RhoA is mediated by AKT phosphorylation of DLC1. J. Cell Biol. 216, 4255–4270 (2017).

30. Ko, F. C. F. et al. PKA-induced dimerization of the RhoGAP DLC1 promotes its inhibition of tumorigenesis and metastasis. Nat. Commun. 4, 1618 (2013).

31. Straszewski-Chavez, S. L. et al. The Isolation and Characterization of a Novel Telomerase Immortalized First Trimester Trophoblast Cell Line, Swan 71. Placenta 30, 939–948 (2009).

32. Yamaguchi, N. & Knaut, H. Focal adhesion-mediated cell anchoring and migration: from *in vitro* to *in vivo*. Development 149, dev200647 (2022).

33. Lauffenburger, D. A. & Horwitz, A. F. Cell Migration: A Physically Integrated Molecular Process. Cell 84, 359–369 (1996).

34. Kouchi, Z. & Kojima, M. Function of SYDE C2-RhoGAP family as signaling hubs for neuronal development deduced by computational analysis. Sci. Rep. 12, 4325 (2022).

35. Coso, O. A. et al. The small GTP-binding proteins Rac1 and Cdc42regulate the activity of the JNK/SAPK signaling pathway. Cell 81, 1137–1146 (1995).

36. Minden, A., Lin, A., Claret, F.-X., Abo, A. & Karin, M. Selective activation of the JNK signaling cascadeand c-Jun transcriptional activity by the small GTPases Rac and Cdc42Hs. Cell 81, 1147–1157 (1995).

37. Teramoto, H. et al. Signaling from the Small GTP-binding Proteins Rac1 and Cdc42 to the c-Jun N-terminal Kinase/Stress-activated Protein Kinase Pathway. J. Biol. Chem. 271, 27225–27228 (1996).

38. Teramoto, H. et al. The Small GTP-binding Protein Rho Activates c-Jun N-terminal Kinases/Stress-activated Protein Kinases in Human Kidney 293T Cells. J. Biol. Chem. 271, 25731–25734 (1996).

39. David, M., Petit, D. & Bertoglio, J. Cell cycle regulation of Rho signaling pathways. Cell Cycle 11, 3003–3010 (2012).

40. Cherfils, J. & Zeghouf, M. Regulation of Small GTPases by GEFs, GAPs, and GDIs. Physiol. Rev. 93, 269–309 (2013).

41. Havrylov, S., Rzhepetskyy, Y., Malinowska, A., Drobot, L. & Redowicz, M. Proteins recruited by SH3 domains of Ruk/CIN85 adaptor identified by LC-MS/MS. Proteome Sci. 7, 21 (2009).

42. Sheftic, S. R., Page, R. & Peti, W. Investigating the human Calcineurin Interaction Network using the πϕLxVP SLiM. Sci. Rep. 6, 38920 (2016).

43. Saso, J., Shields, S.-K., Zuo, Y. & Chakraborty, C. Role of Rho GTPases in Human Trophoblast Migration Induced by IGFBP11. Biol. Reprod. 86, (2012).

44. Ma, X.-L. et al. Upregulation of RND3 Affects Trophoblast Proliferation, Apoptosis, and Migration at the Maternal-Fetal Interface. Front. Cell Dev. Biol. 8, 153 (2020).

45. Ridley, A. J. & Hall, A. The small GTP-binding protein rho regulates the assembly of focal adhesions and actin stress fibers in response to growth factors. Cell 70, 389–399 (1992).

46. Ridley, A. J., Paterson, H. F., Johnston, C. L., Diekmann, D. & Hall, A. The small GTP-binding protein rac regulates growth factor-induced membrane ruffling. Cell 70, 401–410 (1992).

47. Kozma, R., Ahmed, S., Best, A. & Lim, L. The Ras-Related Protein Cdc42Hs and Bradykinin Promote Formation of Peripheral Actin Microspikes and Filopodia in Swiss 3T3 Fibroblasts. Mol. Cell. Biol. 15, 1942–1952 (1995).

48. Nobes, C. D. & Hall, A. Rho, Rac, and Cdc42 GTPases regulate the assembly of multimolecular focal complexes associated with actin stress fibers, lamellipodia, and filopodia. Cell 81, 53–62 (1995).

49. Martin, K. et al. Spatio-temporal co-ordination of RhoA, Rac1 and Cdc42 activation during prototypical edge protrusion and retraction dynamics. Sci. Rep. 6, 21901 (2016).

50. Pertz, O., Hodgson, L., Klemke, R. L. & Hahn, K. M. Spatiotemporal dynamics of RhoA activity in migrating cells. Nature 440, 1069–1072 (2006).

51. Miller, A. L. & Bement, W. M. Regulation of cytokinesis by Rho GTPase flux. Nat. Cell Biol. 11, 71–77 (2009).

52. Hinderling, L. et al. GTPase-activating protein DLC1 spatio-temporally regulates Rho signaling. eLife 12, RP90305 (2026).

53. Kreider-Letterman, G. et al. ARHGAP17 regulates the spatiotemporal activity of Cdc42 at invadopodia. J. Cell Biol. 222, e202207020 (2023).

54. Miller, C. J. et al. Comprehensive profiling of the STE20 kinase family defines features essential for selective substrate targeting and signaling output. PLOS Biol. 17, e2006540 (2019).

55. Arur, S. & Schedl, T. Generation and purification of highly specific antibodies for detecting post-translationally modified proteins in vivo. Nat. Protoc. 9, 375–395 (2014).

56. Torres Robles, J., Stiegler, A. L., Boggon, T. J. & Turk, B. E. Cancer hotspot mutations rewire ERK2 specificity by selective exclusion of docking interactions. J. Biol. Chem. 301, 108348 (2025).

57. Schindelin, J., et al. Fiji: an open-source platform for biological-image analysis. Nat. Methods 9, 676–682 (2012).

58. Horzum, U., Ozdil, B. & Pesen-Okvur, D. Step-by-step quantitative analysis of focal adhesions. MethodsX 1, 56–59 (2014).

59. Tinevez, J.-Y. et al. TrackMate: An open and extensible platform for single-particle tracking. Methods 115, 80–90 (2017).

60. Ershov, D. et al. TrackMate 7: integrating state-of-the-art segmentation algorithms into tracking pipelines. Nat. Methods 19, 829–832 (2022).

