## Supplemental Figure 1 for "Functional characterization of Rho GTPase activating proteins SYDE1 and SYDE2"

Supplemental figure for Song, et al., Functional characterization of Rho GTPase activating proteins SYDE1 and SYDE2

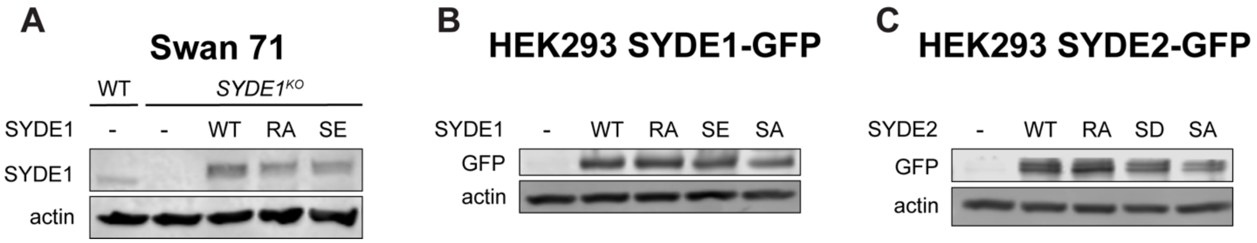


**Fig. S1. Immunoblots showing expression of SYDE protein GFP fusions in Swan 71 and HEK293 cells**

(**A**) Immunoblots of lysates from Swan 71 WT or Swan 71 *SYDE1*$^{KO}$ cells expressing either GFP (-) or the indicated forms of SYDE1-GFP. (**B**) Immunoblots of lysates from HEK293 lysates expressing GFP (-) or the indicated forms of SYDE1-GFP. (**C**) As panel B for cells expressing variant forms of SYDE2-GFP.
